# Gas1-Mediated Suppression of Hepatoblastoma Tumorigenesis

**DOI:** 10.1101/2024.10.02.616350

**Authors:** Keyao Chen, Huabo Wang, Bingwei Ma, Jessica Knapp, Colin Henchy, Jie Lu, Taylor Stevens, Sarangarajan Ranganathan, Edward V. Prochownik

**Author notes:** Correspondence: Edward V. Prochownik, MD, PhD, Division of Hematology/Oncology, UPMC Children’s Hospital of Pittsburgh, Rangos Research Center, Room 5124, 4401 Penn Ave., Pittsburgh, PA 15224. These authors contributed equally to this work (Keyao Chen, Huabo Wang). **AUTHOR CONTRIBUTIONS** Keyao Chen performed experiments, prepared figures and revised the manuscript. Huabo Wang organized and analyzed genomics data. Bingwei Ma, Jessica Knapp, Colin Henchy, Jie Lu and Taylor Stevens performed experiments and collected data. Sarangarajan Ranganathan reviewed and interpreted pathology. Edward V. Prochownik conceived the study, organized the data and wrote the manuscript. All authors reviewed the final version of the manuscript.

## Abstract

**Background and Aims:** Hepatoblastoma (HB), the most common pediatric liver cancer, often dysregulates the Wnt/β-catenin, Hippo and NFE2L2/NRF2 pathways. Pairwise combinations of oncogenically active forms of the terminal transcription factor effectors of these pathways, namely β-catenin (B), YAP (Y) and NRF2 (N) generate HBs in mice, with the triple combination (B+Y+N) being particularly potent. Each tumor group alters the expression of thousands of B-,Y- and N-driven unique and common target genes. Identifying those most responsible for transformation is thus an important question as it might reveal new mechanistic insights and therapeutic opportunities.

**Approach and Results:** Transcriptional profiling of >60 murine HBs driven by the above oncogenic combinations and different B mutants and in genetic backgrounds that impair tumor growth rates but not initiation has revealed a common set of 22 “BYN genes” that are similarly deregulated in all cases. Many are associated with multiple “Cancer Hallmarks” and their expression levels often correlate with survival in human HBs, hepatocellular carcinomas and other cancers. Among the most down-regulated of these is *Gas1*, which encodes a Glycosylphosphatidylinositol (GPI)-linked outer membrane protein. We show here that restoring Gas1 expression impairs B+Y+N-driven HB tumor growth *in vivo* and in HB-derived immortalized cell lines *in vitro* in a manner than requires membrane anchoring of the protein via its GPI moiety.

**Conclusions:** Our findings implicate Gas1 as a proximal mediator of HB pathogenesis and validate the BYN gene set as one deserving of closer additional scrutiny in future studies.

## INTRODUCTION

Hepatoblastoma (HB), the most common pediatric liver cancer, is usually diagnosed in children less than 3-4 years of age.^1^ Believed to originate in developmentally-restricted fetal hepatoblasts, HBs are currently classified into several histologically distinct subtypes despite being the least molecularly complex of all cancers.^1-4^ The most common molecular changes involve the Wnt/β-catenin, NFE2L2/NRF2 and Hippo signaling pathways.^5^ More specifically, they involve missense or in-frame deletion mutations of the *CTNNB1* gene, which encodes β-catenin (B), amplification of or missense mutations in the *NFE2L2/NRF2* (N) gene and aberrant nuclear accumulation of the Hippo terminal effector YAP (Y).^5-8^ This relative molecular simplicity has been reconciled with HB’s histologic and biological diversity by demonstrating that the *in vivo* expression in mice of different pairwise combinations of B, Y and N mutants generates tumors with distinct growth rates, histologies and transcriptional profiles that recapitulate some of the above-mentioned human subtypes.^5-7,9^ Moreover, the triple combination of B, Y and N drives a particularly aggressive form of HB closely resembling the so-called “crowded fetal” histologic subtype and is accentuated by innumerable fluid-filled cysts that are otherwise a rare feature of human HB.^6,10^ Further diversity can be generated by different patient-derived B mutant proteins.^7,9^ Distinct and diverse experimental HB characteristics and behaviors can thus be recurrently generated in mice by over-expressing different combinations of the above factors and B mutants from this otherwise simple oncogenic menu.^5^

B, Y and N are all transcription factors (TFs), with hundreds-thousands of unique and shared direct target genes.^6,7,11-13^ HBs can differ widely in their gene expression profiles in ways that reflect different combinations of the TFs and B mutation identities.^5-7^ This raises the question of which of the these downstream genes are directly responsible for transformation. Among the most highly expressed genes in HBs is *MYC*, a downstream target of B,Y and N that encodes another oncogenic TF.^14,15^ This indicates the existence of a “transcriptional cascade” that amplifies the upstream factors and confers additional gene expression complexity to different HB subtypes. However, unlike B, Y and N, Myc is not mandatory for transformation as HBs can still be efficiently generated from *Myc-/-* murine hepatocytes although tumor growth is slowed.^16^ Further augmenting this transcriptional cascade is the Mlx Network that closely cooperates with the structurally and functionally related Myc Network.^16-18^ The hepatocyte-specific loss of ChREBP, the Myc-like member of this network, whose gene is also a B,Y and N target, also impairs HB growth without affecting initiation as does the individual or combined inactivation of Mlx, Chrebp’s obligate heterodimerization partner.^17^ The ensuing transcriptional heterogeneity among these various molecular classes of HBs further complicates the identification of the most relevant underlying oncogenic initiators.

The availability of the above murine models of HB has previously allowed us to compare their transcriptional profiles. Doing so identified 22 common genes that are invariably up- or downregulated among all murine HBs.^5,6^ 18 of these “BYN genes” are associated with an average of 3.4 “Cancer Hallmarks” versus 6.4 for well-known oncogenes and tumor suppressors (TSs) such as *MYC, RAS, TP53* and *RB*.^19^ Furthermore, the expression patterns of 10 BYN genes could distinguish between human HBs with favorable and unfavorable outcomes, which complemented other studies.^6,11,13^ Finally, 19 BYN genes were dysregulated in murine hepatocellular carcinomas (HCCs) driven by a *MYC* transgene and 17 correlated with survival in over a dozen human cancer types.^6^ Many BYN genes encode enzymes, membrane-associated or extracellular proteins, thereby suggesting that they might be less challenging therapeutic targets than any upstream TF.^20^ Among the more prominent down-regulated genes was *Gas1* (Growth arrest-specific 1), whose transcriptional suppression correlated inversely with the growth rates and aggressiveness of different HB molecular subtypes.^6^

*Gas1* was originally identified as being induced in NIH3T3 murine fibroblasts in response to serum starvation and the only such protein whose enforced re-expression inhibited serum-stimulated G_0_ → G_1_ progression.^21-26^ Mature murine Gas1 is a ∼37 kDa glycosylphosphatidylinositol (GPI)-linked outer membrane protein that exerts pleiotropic effects on survival, apoptosis and mitogenic signaling.^21,26-28^ It is synthesized as a ∼40 kDa precursor that, like other GPI-linked proteins, contains an N-terminal ∼38 amino acid signal peptide that directs Gas1 to the ER where, upon signal peptide removal, an additional ∼28 amino acid C-terminal segment is cleaved and replaced by a serine-linked GPI moiety.^26,29^ Upon further processing by the Golgi apparatus, the mature Gas1 protein, anchored by this GPI tag, is displayed on the outer plasma membrane. However, whether this “mature” post-translationally processed and modified protein represents the only form that mediates cell cycle arrest and other biological functions has been questioned by studies showing that neither C-terminal modification nor GPI addition is necessary for these activities in certain cell types.^26,30,31^

In some cases, Gas1’s suppression of normal and tumor cell growth is at least partially dependent upon TP53 but does not require the latter’s transcriptional activation function. Rather, it requires a proline-rich domain in TP53 located immediately downstream of the N-terminal transactivation domain.^21,32^

We have examined here Gas1’s putative role as a HB TS in the context of its membership in the BYN gene collection. We find *Gas1*’s regulation to be complex but also dependent upon the coordinate activities of the aforementioned TFs that initiate HB tumorigenesis and growth rate. We also show that Gas1 transcript and protein down-regulation is a common feature of many human and murine HBs and HCCs. Finally, we demonstrate that the enforced over-expression of Gas1 impairs murine HB tumorigenesis *in vivo*. A more detailed assessment of Gas1’s effect *in vitro* show that its inhibition of proliferation is dependent upon its proper GPI-dependent display on the outer surface of the plasma membrane. Collectively, our findings support the conclusion that enforcing the expression of a prominent TS-like BYN member can directly alter HB pathogenesis by inhibiting tumor cell proliferation. The findings support the idea that BYN gene members, either individually or collectively, can contribute to HB pathogenesis, are regulated by the major TFs that drive the disease and represent the most proximal factors that contribute to transformation.

## EXPERIMENTAL PROCEDURES

### Animal care and husbandry

Four-six week old FVB/N mice were purchased from Jackson Labs, Inc. (Bar Harbor, ME) and were used for all *in vivo* tumor studies. They were maintained in micro-isolator cages with *ad libitum* access to standard mouse chow and water. All care and procedures were reviewed and approved by The University of Pittsburgh’s Department of Laboratory and Animal Resources (DLAR) and the Institutional Animal Care and Use Committee (IACUC).

### Bacterial plasmids

Sleeping Beauty (SB) vectors encoding the Δ90 patient-derived in-frame B deletion mutation, Y^S127A^, N^L30P^ and a non-SB vector encoding SB transposase (pCMV-transposase) have been previously described.^7,17,18^ The same SB vector encoding a V5 epitope-tagged full-length murine Gas1 protein was synthesized by GenScript Biotech (Piscataway, NJ) and was used as a template for the re-cloning of the cDNA into another SB vector, pSBbi-RP (Addgene, Inc., Watertown, MA). This was utilized for the expression of the original full-length (343 amino acid) unprocessed murine Gas1 protein (Uniprot ID Q01721) that included the N-terminal ER targeting sequence, the C-terminal moiety that must be cleaved prior to GPI addition and a C-terminal V5 epitope tag. The absence of V5 epitope-tagged Gas1 protein generated by this vector was used to confirm the efficiency of post-translational cleavage of the C-terminus in the fully processed protein. Finally, a second pSBbi-RP vector encoding a truncated Gas1 protein, lacking the C-terminal 29 amino acids (and the V5 tag) and predicted to be secreted due to its inability to be anchored to the plasma membrane via its GPI modification, was used to determine whether soluble Gas1 inhibited HB cell line growth. cDNAs encoding both of these proteins were amplified from the original SB vector using the common forward primer 5’-TCA AGC CTC AGA CAG TGG TTC-3’. The sequence of the reverse PCR primer used to amplify the full-length cDNA was: 5’-TAG AAG GCA CAG TCG AGG-3’ and the sequence used to amplify the 29 codon-truncated cDNA was: 5’-ACT GTC TAG ATT AGC CGC TAG AGG AGC TCC GT-3’. All oligonucleotides were synthesized by IDT, Inc. (Coralville, IA). After digesting with NcoI and XbaI, cDNAs were cloned directionally into the pSBbi-RP vector that contained these same sites.

Two pDG458 Crispr/spCas9 vectors (pDG1+2 and pDG3+4)^33^ were used to express 4 gRNAs against exon 2 of the murine *Cdkn2a* gene. The top strand sequences were: Oligo 1: 5’-CGGTGCAGATTCGAACTGCGAGG-3’; Oligo 2: 5’-GTCGTGCACCGGGCGGGAGAAGG-3’; Oligo 3: 5’-CTTGGGCCAAGTCGAGCGGCAGG-3’ and Oligo 4: TGCGATATTTGCGTTCCGCTGGG-3’. Another pDG458 vector was generated that encoded 2 gRNAs (pDG5+6) against exon 1α. This was used to specifically inactivate p16^INK4A^ while preserving p19^ARF^.^34^ Top strand sequences for this vector were: Oligo 5: 5’-AGGGCCGTGTGCATGACGTGCGG-3’ and Oligo 6: 5’ CTCCTTGCCTACCTGAATCGGGG-3’. Each double-stranded oligonucleotide was inserted into its respective site in pDG458 Crispr/spCas9 using a previously described single-step digestion-ligation procedure (https://media.addgene.org/data/plasmids/100/100900/100900-attachment_Yl0i43bWJig3.pdf). After confirming all sequences by di-deoxy DNA sequencing, plasmids were purified using Plasmid Plus Midi columns (Qiagen, Inc., Germantown, MD), re-dissolved in sterile TE buffer and stored at -20°C.

### Generation of HBs

HBs were induced in 6-8 wk old mice as previously described.^7,17,18^ In brief, purified plasmids were delivered to the liver by hydrodynamic tail vein injection (HDTVI) in 2 ml of PBS over 5-10 sec. Each inoculum included 10 μg each of the indicated SB vectors along with 2 μg of the pCMV transposase-encoding vector. Tumors with *Cdkn2a* locus mutations were generated in the same manner except that 2 μg of each pDG458 vector were included in each inoculum.

### Murine HB cell lines

The BY1 and BY2 murine HB cell lines were generated from primary BY tumors in which the *Cdkn2a* locus was disrupted by targeting exon 2 using the pDG458(1+2) and pDG458(3+4) Crispr/Cas9 vectors described above.^35^ A third cell line, BY21, was generated by targeting *Cdkn2a* exon 1α using the pDG458(5+6) vector only. Upon reaching maximal size, BY tumors carrying these mutations were excised. Approximately 2 g of tumor was minced, washed several times in PBS and digested for 30 min at 37°C in 0.1% trypsin (Sigma-Aldrich, Inc. St Louis, MO). After further disruption by vigorous pipetting and vortexing, the digested samples were added to tissue culture plates containing standard D-MEM-10% FBS. Over the next 1-2 wks, macroscopic remnants of the original tumor tissue were discarded. After several additional weeks, both microscopically and macroscopically visible colonies could be identified amidst the background of non-proliferating cells. These were further expanded and then frozen in liquid nitrogen within 3-5 additional passages.^35^ BY1, BY2 and BY21 cell lines were transfected with 2 μg the above-mentioned pSBbi-RP-Gas1 expression vectors and 0.2 μg the pCMV-transposase vector using Lipofectamine-3000 according to the directions of the vendor (Thermo-Fisher, Inc., Waltham, MA). Control cell lines were established from cells transfected with the empty pSBbi-RP vector alone. All transfected cells were selected in puromycin (2 μg/mL) and expressed dTomato that was also encoded by the pSBbi-RP vector. In other experiments, transiently transfected cells were separated by fluorescence-activated cell sorting based on the intensity of dTomato expression. All cell sorting utilized a BD FACSAria II Instrument (Becton-Dickinson, Inc. Franklin Lakes, NJ). Cell counting was performed on an Incucyte Live-Cell Imaging and Analysis System (Sartorius Essen, Inc. Ann Arbor, MI) as previously described.^35^

### Immunoblotting

Tissue fragments and cell pellets were lysed in 1 X SDS-PAGE buffer in the presence of protease and phosphatase inhibitors.^7,16,17^ Following standard denaturing SDS-PAGE, separated proteins were blotted to Millipore PVDF membranes (Burlington, MA) using a semi-dry apparatus (Fisher™ Semi-dry Blotting Apparatus, Thermo Fisher). Antibodies used for immunoblotting and immunofluorescence studies included those for Gas1 (1:1000, R&D Systems, Inc. Minneapolis, MN, cat. no. AF2644-SP), V5 epitope tag (1:5000, Thermo Fisher, Inc., Waltham, MA, 1:5000), GAPDH (1:5000, Sigma-Aldrich, Inc. St. Louis, MO, Cat. no. G8795), Akt (1:500, Santa Cruz Biotechnology, Inc. Dallas, TX, cat. no. sc-5298), pSer^473^ Akt (1:500, Cell Signaling Technologies [CST], Inc., Danvers, MA, cat. no. 9271), Actin (1:5000, CST, cat. no. 3700), HRP-conjugated goat anti-mouse IgG (1:5000, CST) and HRP-conjugated goat anti-rabbit IgG (1:5000, CST). Blots were developed using a chemiluminescence kit (Thermo Fisher, cat. no. ThemoFisher 34096) and Imaged on Protein Simple, Fluoro Chem M System (Bio-Techne, Inc., Minneapolis, MN).

### Computational analyses

Consensus TF binding sites within the intron-less *Gas1* gene were retrieved from the JASPAR database through the University of California Santa Cruz Genome Browser. The sites identified included those for β-catenin (Tcf4 sites), Yap/Taz (TEAD sites), Myc (E boxes), Nrf2 (ARE sites), Sp1 and Miz1, each with scores ranging from ∼300-370. ChIP-seq data, sourced from ReMap, were obtained through Genome Browser. Representative ChIP-seq peak maps were either downloaded from the Gene Expression Omnibus (GEO) database or subjected to reanalysis using the provided raw sequencing data. A schematic cartoon depicting the binding sites and CHIP-seq peaks was generated using R version 4.3.1, with the ggplot2 package employed for visualization.

To quantify *Gas1* and other transcript levels in murine HBs induced by various B,Y and N combinations and in murine HCCs induced by conditional Myc over-expression, we queried previously published RNAseq results from the GEO database.^7,16,17,36^ Quantification of *Gas1* transcripts in select human cancers from The Cancer Genome Atlas (TCGA) was performed as described previously.^6^ Briefly, RNAseq expression data (FPKM-UQ) and clinical annotation files were downloaded from the TCGA GDC PANCAN dataset and accessed through Xenabrowser. Patients were divided into high-*Gas1* and low-*Gas1* expression groups using a cutoff that maximized survival differences. Survival analysis for each group was performed using R version 4.3.1 with the package Survival.

## Statistical analysis

GraphPad Prism 8.0 was used for all analyses and employed a 2-tailed unpaired Student’s t test or a Mann-Whitney U test for comparing differences between 2 groups. For comparison among more than 2 groups, one-way ANOVA with multiple comparison were used and adjusted p-value was reported. Data are shown as means ± SEM. Kaplan-Meier survival curves were generated using the log-rank test. At least 3 biological replicates were performed for each experiment.

## RESULTS

### Simultaneous binding of B,Y, N and Myc to sites in the human and mouse *Gas1* genes suggests interactive and conserved transcriptional regulation

The intron-less mouse and human *Gas1* genes contain multiple and seemingly evolutionarily conserved consensus binding sites for β-catenin (Tcf4 sites), YAP/TEAD (TEAD sites), Nrf2 (ARE sites), Myc (E boxes) and Sp1 and Miz1 (at their eponymous binding sites) (Figures 1A, B). Sp1 and Miz1 are positively-acting TFs that are directly inhibited by Myc and regulate the majority of negative Myc targets.^16,37,38^ ChIP seq results from ENCODE showed that each of the above TFs directly bound both the human and murine *Gas1* genes in proximity to their transcriptional start and end sites (TSS and TES, respectively) and, in most cases, within the genes’ coding regions as well (Figures 1C, F).^39^ Myc binding often overlapped Sp1 DNA elements while also coinciding with sites of actual Sp1 occupancy. This was particularly prominent around the TSS where Sp1 binding also overlapped that of other factors. Elsewhere, binding sites aligned imperfectly with their respective DNA binding elements in the gene body. For example, despite the presence of prominent NRF2 footprints, none were associated with ARE binding sites in either the human or mouse genes. This implicated NRF2 binding as being indirect and perhaps occurring in association with other factors such as Myc or at non-consensus DNA elements. The loose coincidence of Myc and NRF2 binding sites supported the former possibility and was in keeping with their previously described interaction.^40^ Collectively, our findings revealed *Gas1* gene regulation to be complex, evolutionarily conserved and responsive to numerous TFs relevant to HB causation, both individually and combinatorially.

**Figure 1.**
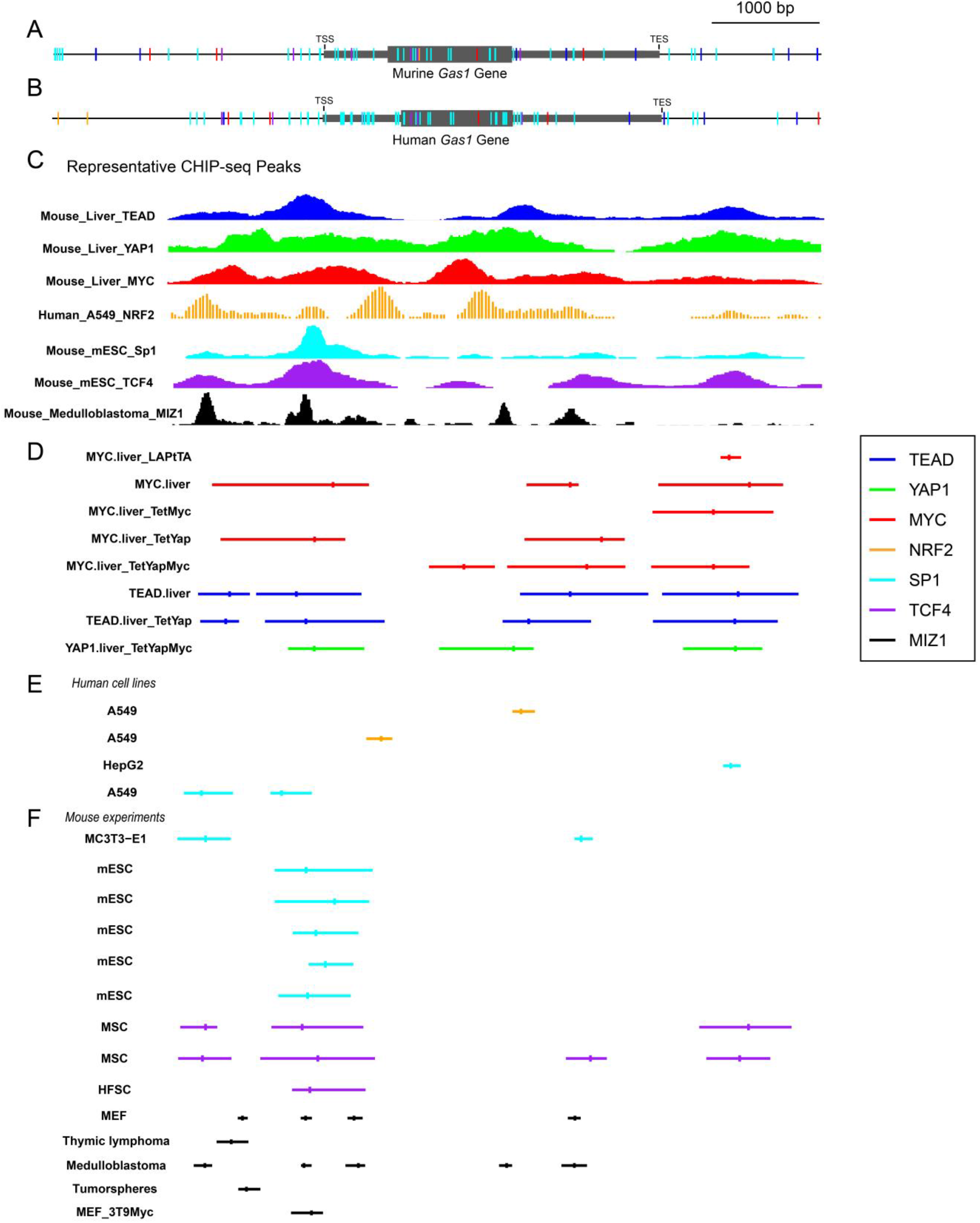
Conserved binding of relevant TFs to the murine and human *Gas1* genes. (A,B) Cartoons of the intron-less murine *Gas1* and human *Gas1* genes, respectively, showing locations of consensus and evolutionarily conserved binding sites for the indicated relevant TFs. TSS: transcriptional start site; TES: transcriptional end site. (C) Representative examples of binding footprints from the following experiments shown in panel D,E,F for the indicated TFs in relation to the binding sites shown in A and B from 4 different cell types/tissues (murine liver, human A549 lung cancer cells, murine embryonic stem cells [mESCs] and murine medulloblastoma). ^39^ (D) Summaries of all available ENCODE data for *Gas1* gene binding by Myc^39^ and YAP/TEAD from mouse livers, mouse livers over-expressing YAP and Myc-driven HCCs [GSE83863],. Hash marks indicate the geometric mean of TF binding distribution across different experiments. (E) Summaries of ENCODE data for *Gas1* gene binding by NRF2 and Sp1 from A549 lung cancer [GSE141497, GSE113497, ENCSR000BPE], and HepG2 HB cell lines [ENCSR000BJX].^39^ (F) Summaries of ENCODE data for *Gas1* gene binding of Sp1, Tcf4 and Miz1 in murine MC3T3 cells [GSE76185], mESCs [GSE126496], mesenchymal stem cells (MSC) [GSE137089], murine embryonic fibroblasts (MEF), hair follicle stem cells (HFSC) [GSE48878], medulloblastoma tumor spheres [GSM1571163] and Myc over-expressing 3T9 fibroblasts [GSE98419].

### Gas1 transcripts and protein are down-regulated in liver cancer

*Gas1* transcripts were markedly lower in murine HBs driven by each combination of B, Y and N. This was most prominent in B+Y+N tumors, the most aggressive of the 4 subtypes (Figure 2A).^6^ HBs generated by a combination of Y and other in-frame deletion and missense mutations of B showed a similar depression of *Gas1* transcripts as did slowly growing B+Y tumors generated in *Myc-/-, Chrebp-/-* and (*Myc-/-*x*Chrebp-/-*) “double knockout” hepatocyte backgrounds (Figure 2B).^6,16,17^ Consistent with these findings, analysis of two previously reported RNAseq studies of 59 primary human HBs and matched livers showed *Gas1* transcript levels to be significantly lower in tumors with higher molecular risk scores (Figure 2C).^12,13^

**Figure 2.**
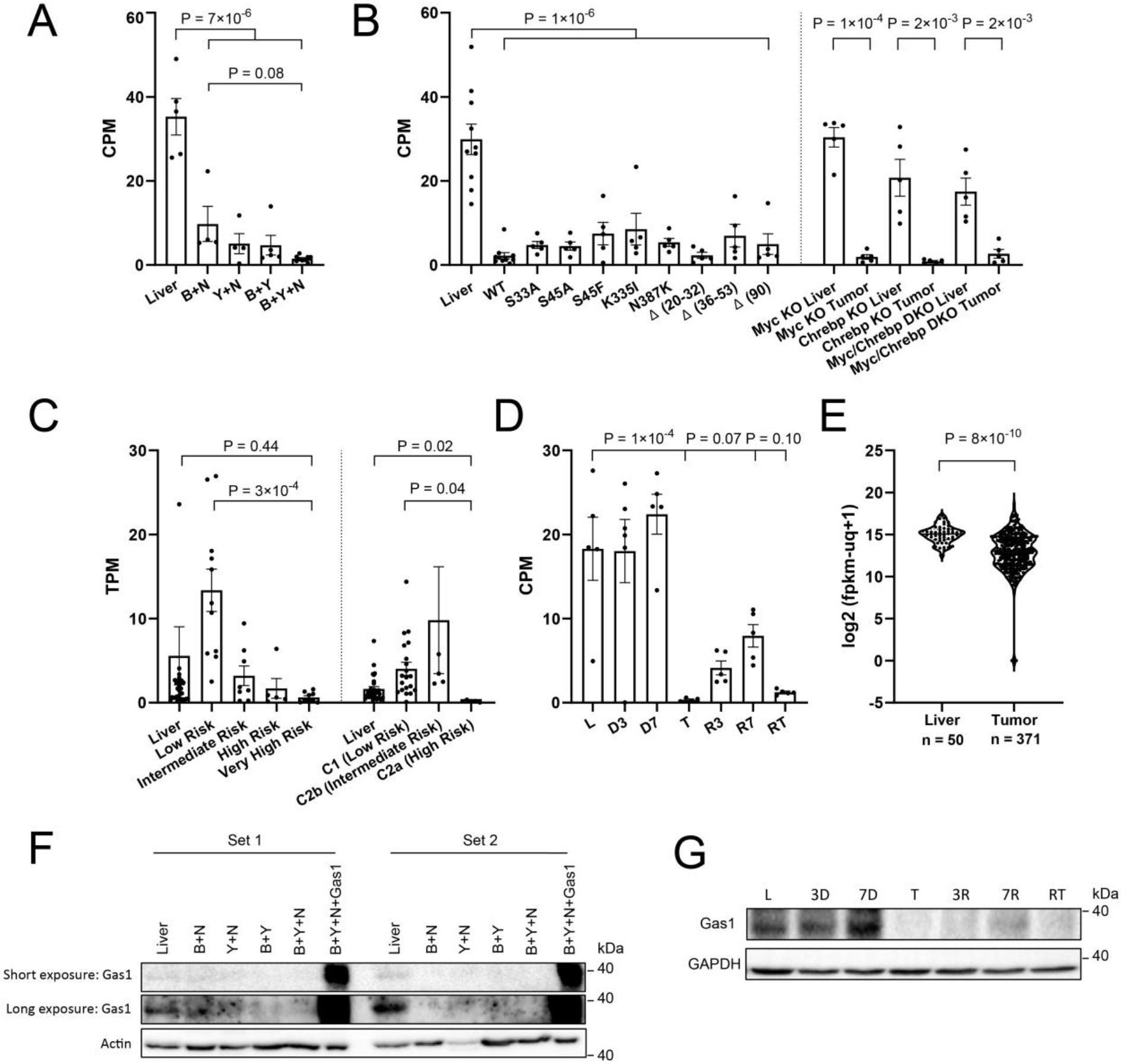
Gas1 transcript and protein levels decline in liver cancers. (A) Expression of *Gas1* transcripts in livers and murine HBs generated by the indicated combinations of B,Y and N and obtained from RNAseq profiling.^6^ (B) *Gas1* transcript levels in murine HBs generated by Y and each of the indicated β-catenin mutants.^7^ Also shown are *Gas1* transcript levels in BY HBs generated in mice with hepatocyte-specific knockouts of *Myc, Chrebp* and *Myc* x *Chrebp* (DKO) (C) *Gas1* transcripts levels in 59 human HBs previously stratified into different risk groups based on prior molecular profiling.^12,13^ (D) *Gas1* transcripts in control livers and HCCs generated by doxycycline-regulated over-expression of a human *MYC* transgene. D3 and D7: 3 days and 7 days, respectively, after *MYC* induction but prior to tumor development^36^; T: tumors obtained ∼30 days after *MYC* induction: R3 and R7: regressing HCCs sampled 3 days and 7 days after silencing *MYC*. RT: recurrent HCC induced 3-4 months after the original tumor’s complete regression. (E) *Gas1* transcripts in human HCCs and adjacent matched livers from the TCGA database. (F) Gas1 protein levels in 2 sets of normal livers and HBs driven by the indicated oncoprotein combinations. The last lane of each set shows the expression of Gas1 in B+Y+N tumors in mice that were also injected with a SB vector encoding Gas1. (G) Gas1 protein expression in the liver and HCC tissues from D.

The induction of undifferentiated HCCs in response to doxycycline-regulated over-expression of a human *MYC* transgene in mice was associated with similarly dramatic declines in *Gas1* transcripts (Figure 2D).^36^ This occurred in tumors but not in livers as late as 7 days after *MYC* induction. Regressing tumors after *MYC* silencing showed partial normalization of *Gas1* expression by day 7. *Gas1* transcript suppression was also seen in recurrent tumors induced 3-4 months after the original ones had completely regressed. *Gas1* transcripts were also downregulated in a large subset of human HCCs from TCGA but did not correlate with survival (Figure 2E and not shown). The greater variability of *Gas1* expression among these tumors compared to *MYC*-induced murine HCCs or HBs may reflect their greater molecular complexity or their more variable etiologies and accompanying pathologies.^2,3,5,9,41^

Gas1 protein expression in murine HBs tended to mirror that of *Gas1* transcripts (Figure 2F). B+Y HBs stably expressing a SB-Gas1 vector demonstrated 5-10-fold higher levels of Gas1 than control livers. In both cases, the protein’s sizes were identical, indicating that the V5 epitope tag of the exogenous protein was efficiently removed during post-translational processing.^26,42^

Endogenous Gas1 protein expression in *MYC*-induced murine HCCs also reflected that of transcripts at each time point (Figure 2G). Thus, declines in Gas1 protein were not appreciated until tumors had developed around day 30. Small increases in Gas1 protein expression during the early induction phase (D3-D7) suggested that this might represent an abortive attempt by incipient tumor to restrain uncontrolled growth. As observed with *Gas1* transcripts in regressing tumors, protein levels partially normalized by day 7 and then again became undetectable in recurrent tumors.

### Gas1 inhibits murine HB growth

To determine whether Gas1 functions as a TS *in vivo* as hypothesized, we compared the survival of mice bearing B+Y+N+Gas1 HBs to those bearing control B+Y+N tumors. The former group showed ∼50% longer survival and their tumors expressed high levels of Gas1 (Figure 3A, B).

**Figure 3.**
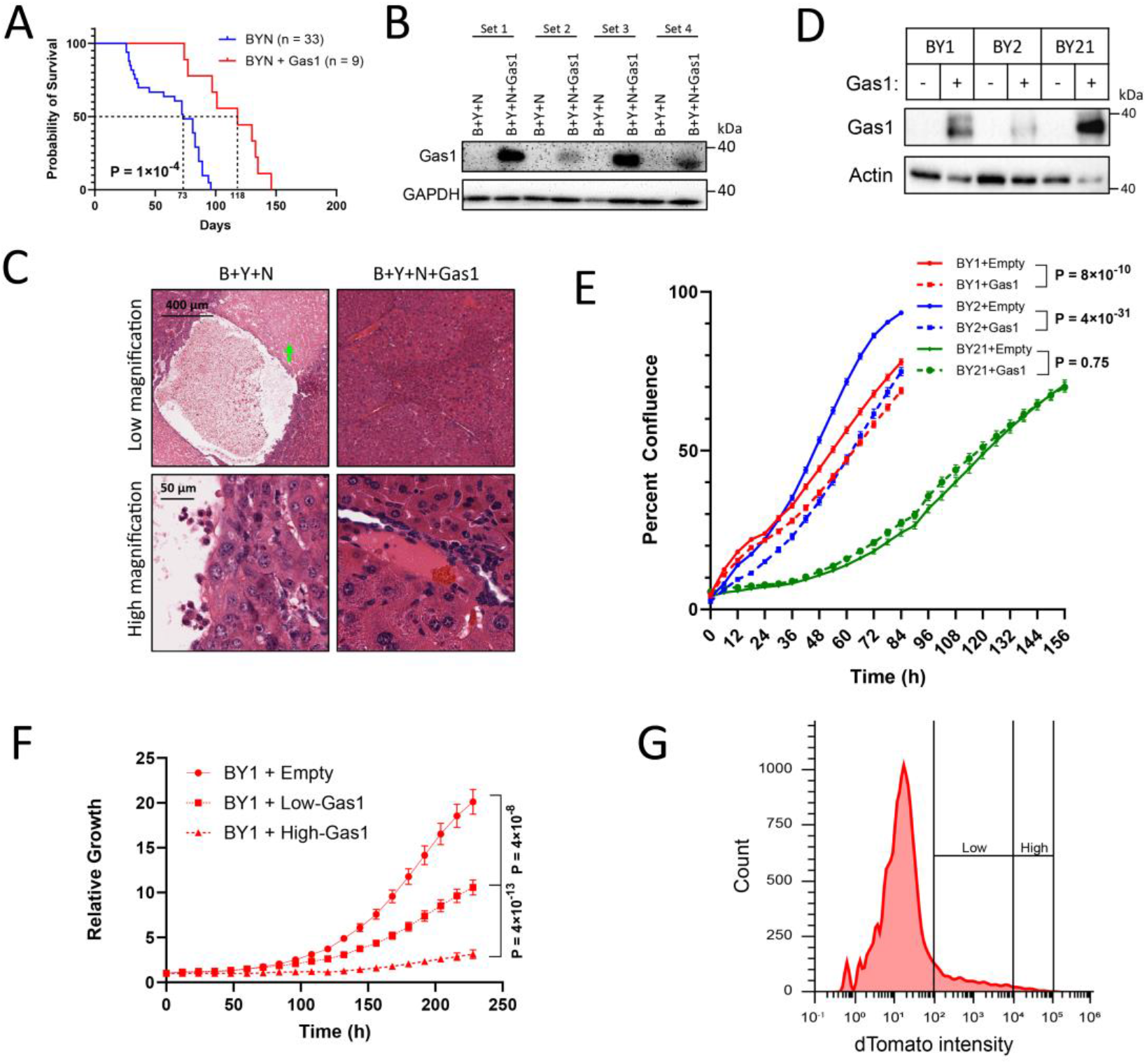
Gas1 restoration extends survival in HB-bearing mice and inhibits tumor cell growth. (A) Kaplan-Meier survival curves. HBs were generated by HDTVI-mediated hepatic delivery of SB vectors encoding B+Y+N or B+Y+N+Gas1.^6,7^ (B) Expression of Gas1 protein in 4 sets of tumors from A. See Figure 2F for additional examples. (C) H&E staining of representative tumors from A. Note fluid-filed cysts with adjacent areas of necrosis (green arrow) in B+Y+N HBs and their absence in B+Y+N+Gas1 HBs. Higher power magnification showed similar tumor cell morphologies resembling the crowded fetal variant of HB.^1^ See for additional examples. (D) Expression of Gas1 in 3 independently derived B+Y cell lines. Each cell line was stably transfected with an empty pSBbi-RP vector or the pSBbi-RP-Gas1-V5 expression vector and selected in puromycin. (E) Gas1 over-expression minimally inhibits BY cell line proliferation. The BY1, BY2 and BY21 cell lines were transfected with a SB-Gas1 expression vector that also encoded dTomato. Two days after transfection, pooled transfected clones were seeded at equal numbers, maintained in puromycin and enumerated at the indicated time points. (F) HB cell growth is inhibited by Gas1 in a dose-dependent manner. Two days after stable transfection with SB-control or SB-Gas1-V5 vectors, cells were sorted into populations with “low” and “high” dTomato expression. Cell numbers were then monitored as in E. (G) Flow diagram of transiently transfected BY1 cells indicating the populations that were designated as expressing “high” and “low” levels of dTomato for the growth studies depicted in F.

B+Y and B+Y+N murine HBs resemble the crowded fetal sub-type of human HB.^1,6,7,43^ B+Y+N HBs however, also contain innumerable fluid-filled cysts, which are rarely observed in human HBs.^6,10^ This was again seen in the control B+Y+N tumors generated for the current study (Figure 3C and Supplemental Figure 1). Also as previously reported, these cysts were often adjacent to well-delineated regions of necrosis.^6^ B+Y+N+Gas1 tumors contained neither of these features thus suggesting that they were either properties of the high growth rates of B+Y+N HBs or that they were suppressed by Gas1.

To evaluate the consequences Gas1 restoration in greater detail, we generated 3 immortalized HB cell lines (BY1, BY2 and BY21) from primary B+Y tumors.^35^ This was enabled by the *in vivo* Crispr-mediated targeting of the *Cdkn2a* gene that inactivated its two encoded TS proteins (p16^INK4A^ and p19^ARF^) (in the case of BY1 and BY2) or p16^INK4A^ only (in the case of BY21).^34,44^ Like the primary tumors from which they originated, these cell lines expressed little-no endogenous Gas1 but did express high levels of the exogenous protein (Figure 3D). Yet, in no case did this inhibit growth by more than ∼15% (Figure 3E). We did note however that the intensity of dTomato expression declined in Gas1-transfected cells suggesting that Gas1 was in fact exerting more growth-suppression than initially supposed. We therefore repeated the above transfections and growth curves with BY1 cells after first sorting high- and low-intensity dTomato-positive populations into separate wells. Over this extended period, we found that dTomato “low” cells grew at about half the rate as control cells whereas dTomato “high” cells grew at only about 1/8^th^ the rate of control cells (Figure 3F-G). We thus conclude that high-level Gas1 expression is selected against but that that maintaning it efficiently suppresses tumor cell proliferation in a manner that is both p16^INK4a^ and p19^ARF^-independent.

### HB growth suppression is dependent upon Gas1’s GPI-dependent attachment to the outer plasma membrane

Growth inhibition by Gas1 is pleiotropic by virtue of its cell surface interactions with glial cell line-derived neurotrophic factor (GDNF) and/or sonic hedgehog (Shh).^26,28,42,45,46^ It also suppresses the growth-promoting PI3-kinase (PI3K)-Akt axis although this is likely related to the latter’s transmission of glial-specific GDNF signaling.^28,47,48^ While Gas1’s effects on these pathways are thought to be facilitated by its GPI-mediated anchoring to the outer membrane, soluble Gas1, lacking the GPI anchor, also acts in a paracrine and/or autocrine manner in association with GDNF, and the co-receptors GFRα1 to repress RET signaling via PI3K-Akt.^23,30,49,50^ To examine this in HB cells, we compared the expression, subcellular localization and growth inhibitory properties of full-length Gas1 and an attachment-defective and secretable form lacking the GPI-modification (Figure 4A). Both proteins were expressed at similar levels within cells with only truncated Gas1 being detected extracellularly (Figure 4B). However, its apparent molecular weight now corresponded to that of the mature GPI-linked wild-type protein seen in whole cell lysates. This suggested that the absence of a GPI tag led to aberrant, non-GPI-associated post-translational modification(s) prior to secretion. Further examination of stably transfected cells by immunofluorescent staining showed mature, GPI-linked protein to be both cytoplasmic- and membrane-localized whereas the GPI-attachment defective form was less abundant and absent from the membrane (Figure 4C). Unlike the growth inhibition imposed by constitutively-expressed mature Gas1 (Figure 3F), GPI attachment-defective Gas1 inhibited growth minimally, if at all (Figure 4D). Tissue culture fluid from these latter cells, even when concentrated >10-folod, was unable to inhibit control BY1 cell growth (Figure 5E). Moreover, no differences in either total or phosphorylated Akt were seen between control cell and those over-expressing either form of Gas1. Finally, based on previously reported RNA-seq data,^6,16,17^ virtually no RET or GDNF transcripts were detected in livers or in BY and BYN HBs (range: 0-0.02 RPKM). HB growth-suppression by Gas1 thus requires its anchoring to the membrane via its GPI moiety and does not mediate its effects via suppression of the GDNF-GFRα1-RET-PI3K-Akt signaling axis.

**Figure 4.**
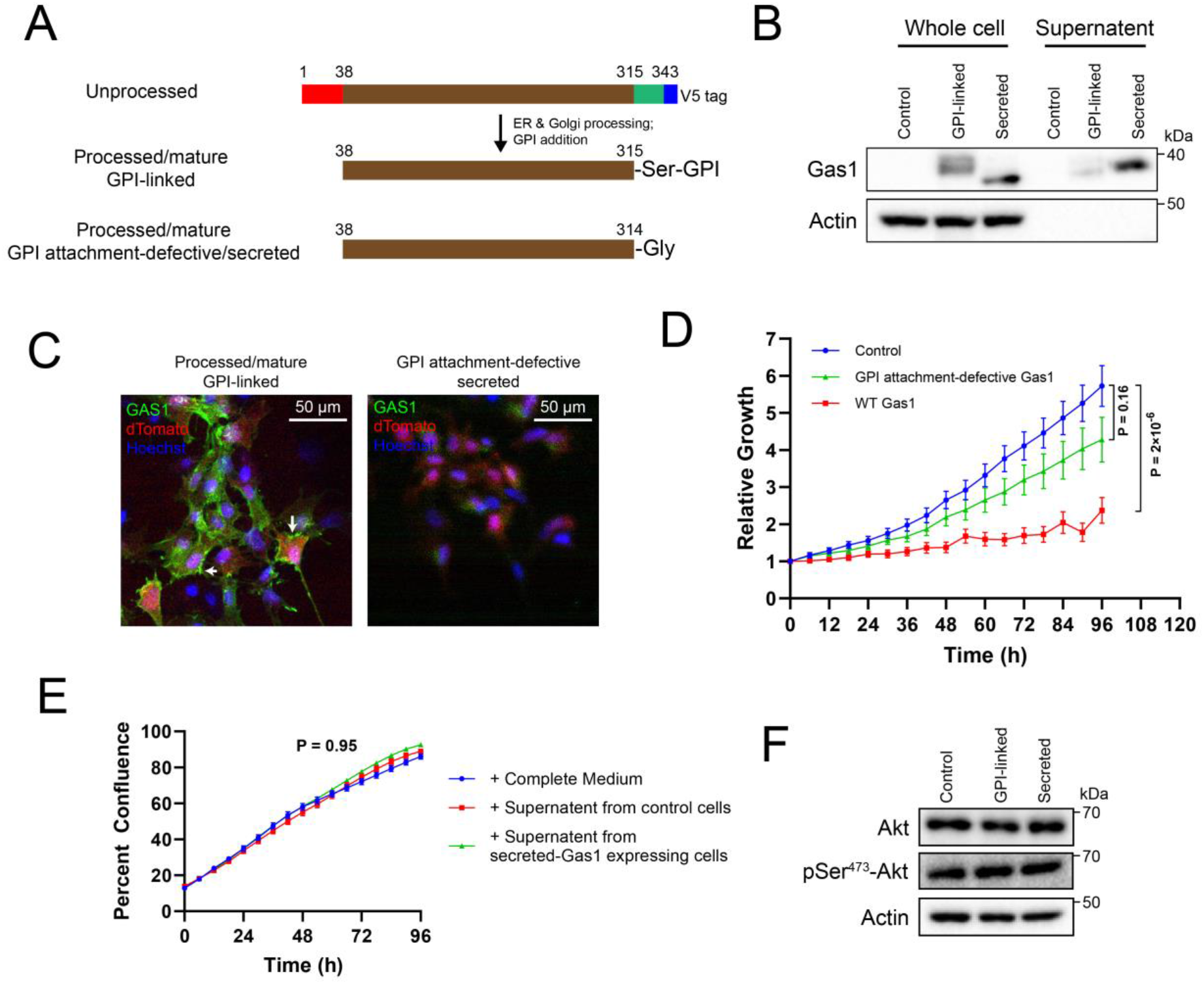
Gas1-mediated growth suppression requires its GPI-mediated tethering to the outer plasma membrane. (A) Structure of Gas1 proteins. Shown at the top is the unmodified, full-length Gas1 precursor protein encoded by the pSBbi-RP vector. In the ER and Golgi, it is converted to the “processed/mature, GPI-linked” form following removal of its N-terminal ∼38 residues (red), its C-terminal ∼28 residues (green) and the 14 residue V5 epitope (blue). It is then further modified by the addition of a GPI moiety to a newly exposed C-terminal serine. The bottom of the cartoon shows the mature version of the GPI-attachment-defective, secreted form of Gas1, which terminates at Gly_314_, and cannot be modified by GPI addition. (B) Expression of the mature forms of Gas1 in transiently transfected BY1 cells. (C) Confocal immunofluorescent images of Gas1 localization (green) in BY1 cells expressing the Gas1 proteins shown in A. White arrows in the first panel indicate regions of membrane localization. dTomato expression (red) is superimposed as is nuclear staining with Hoechst 33342 (blue). In the second panel, cells expressing the GPI attachment-defective secreted form of Gas1 show much less intense Gas1 staining, which is only cytoplasmic. (D) Growth curves of control BY1 cells and those expressing the GPI attachment-defective secreted form of Gas1. The brightest populations of dTomato+ cells were selected and plated as described in Figure 3F and cell proliferation was quantified over a 4 day period. (E) Lack of BY1 cell inhibition by soluble Gas1. Cells expressing the truncated and secreted form of Gas1 (Figure 4A-D) were grown to 100% confluence. After reducing the serum concentration to 0.1%, the supernatant was collected over 3 days. Untransfected naïve BY1 cells were then exposed to the collected medium after restoring the serum concentration to 10%. Cell counts were performed as in D. (F) AKT phosphorylation is not impacted by the over-expression of either mature GPI-linked or GPI attachment-defective Gas1. Control BY1 cells or those over-expressing the indicated Gas1 proteins were subjected to immuno-blotting for total and pSer^473^ Akt.

## DISCUSSION

B,Y and N are TFs with hundreds-thousands of individual and shared positive and negative targets.^6-8,17,36,41,43^ The identification of BYN genes thus represented an attempt to distill the transcriptomic complexity of HBs in order to identify and prioritize the most relevant and proximal drivers of tumorigenesis. An underlying premise of this approach is that such genes are regulated in the same direction regardless of the TF combinations driving the tumor. This regulation should also be evolutionarily conserved and independent of the tumor’s growth rate, its genetic background and the direction of expression of other transcripts without direct roles in transformation.^5,6^ By comparing the gene expression profiles of multiple individual murine HBs generated by all possible combinations of B,Y, and N, by different B mutants and in genetic backgrounds lacking *Myc-/-* and/or *Chrebp-/-*, we identified the 22 member BYN gene set that met these criteria.^6,7,16,17^ Gratifyingly, most of its members possess known functions that could potentially contribute to one or more “Cancer Hallmarks”.^6,19^ Many of them have also been previously appreciated as influencing normal and/or neoplastic proliferation in addition to being deregulated in HCC and other cancers in ways that often correlate with survival.^5,6,11,13^

Among the most highly down-regulated BYN genes (>22-fold in BYN HBs),^6^ Gas1 exerts growth suppressive effects in a variety of both transformed and untransformed cell types.^21,24,26,28,30,31,42,49^ The finding that the functional reversal of this down-regulation, achieved by re-expressing it both *in vivo* and *in vitro*, inhibits tumor cell growth (Figure 3A, F) not only confirms this role in the context of HB but also validates the argument that BYN genes are relevant for tumor initiation as originally proposed and discussed above.^5,6^

Gas1 is invariably suppressed in different molecular subtypes of HB and in Myc-driven HCCs to a degree that reflects the underlying oncogenic driver(s) (Figure 2A-D).^6^ This agrees with the observation that multiple, evolutionarily-conserved binding motifs for β-catenin/Tcf, YAP/Taz, NRF2, Myc, Sp1 and Miz exist within the intron-less *GAS1* gene, with many of these sites being occupied by their respective factors in livers, liver cancers and other normal and neoplastic cell types (Figure 1). This suggests that each factor exerts substantial negative transcriptional control over Gas1 and that cooperative and more nuanced suppression can be achieved combinatorially. This was well-exemplified in HCCs where Myc over-expression alone was sufficient to achieve a degree of Gas1 suppression rivaling that seen in any of the HB groups (Figure 2A, B and D). Notably, most of these factors, as well as the Myc-interacting co-factors Sp1 and Miz1^37,38^ also bound at the gene’s 3’-end in proximity to the TES (Figure 1). We have recently reported that many human and murine genes show Myc- and/or Max-binding in proximity to these sites.^51^ This occurs in association with numerous other TF, co-factors and histone modifiers, an open chromatin environment and local nuclease susceptibility.^51^ Functionally, Myc binding around TESs contributes to chromatin looping while regulating total as well “read-through” transcription that is proposed to modulate nuclear→cytoplasmic transport and translation in response to various stresses, including oncogene over-expression.^51,52^

The immortalized HB cell lines, which were used for much of the current work, could only be generated following concurrent *Cdkn2a* locus inactivation.^35^ *Cdkn2a* encodes 2 critical regulators of proliferation and survival, with p16^INK4a^ promoting replicative and oncogene-induced senescence via the retinoblastoma (Rb) TS pathway and p19^ARF^ indirectly stabilizing the TP53 TS to promote cell cycle arrest and apoptosis.^44,53^ These pathways share considerable cross-talk, with mutations or blocks in each being extremely common in human cancers and immortalized cell lines.^34,54,55^ Gas1 has been reported to rely upon TP53 to promote growth suppression in several cell types.^27,32^ This may explain some of Gas1’s growth-suppressive effects on primary HBs, which contain no Cdkn2a locus mutations and express elevated levels of wild-type p16^INK4A^ and p19^ARF^ (Figure 3A).^35^ In contrast, the immortalized HB cell lines derived from these tumors contain numerous *Cdkn2a* mutations and express no wild-type p16^INK4a^ or p19^ARF^ proteins.^35^ Thus, the growth inhibition that is mediated by mature GPI-linked Gas1 in primary HBs, and that cooperates with *Cdkn2a*-encoded p16^INK4A^ and p19^ARF^, may rely on pathways that are *Cdkn2a-, Rb- and TP53-*independent in HB cell lines. It is thus possible that the lack of any discernible growth inhibition when HB cell lines are cultured in the presence of exogenous extracellular GPI detachment-defective Gas1 (Figure 4E) reflects, at least in part, the *Cdkn2a* differences between primary HB tumors and cell lines.

Soluble, secreted forms of Gas1 lacking the GPI anchor have been reported to inhibit the growth of certain tumor cells, particularly those of neural origin.^29-31^ This occurs via the engagement of Gas1 with the cell surface protein RET, a co-receptor for GDNF, which leads to the suppression of RET-mediated PI3-kinase-Akt signaling.^47-49^ Our inability to affect HB cell growth *in vitro* with high concentrations of GPI attachment-defective secreted Gas1 or to demonstrate any changes in Akt signaling in Gas1-producing cells (Figure 4E, F) likely reflects the tissue differences between these cells and those of neural origin as well as the fact that HBs express little if any RET or GDNF. The fact that mature, GPI-linked Gas1 does seem to markedly inhibit HB cell proliferation both *in vivo* and *in vitro* suggests that it may instead function via an alternate pathway involving inhibitory interactions with the hedgehog pathway leading to cell cycle arrest.^46^ This, plus GPI- and *Cdkn2a*-independent and cytoplasmically-localized Gas1 function (Figure 4C) appear to be the major means by which growth inhibition of HB cell lines is achieved and maintained.

Importantly, the ectopic restoration of Gas1 slows but does not prevent HB tumorigenesis driven by the triple combination of B+Y+N that is associated with particularly aggressive HBs and robust Gas1 suppression (Figure 3A).^6^ Thus, the current work, while indicating that Gas1, does function to inhibit tumor growth as originally postulated, also indicates that other factors, quite likely other members of the BYN gene group must be concurrently normalized to achieve a more durable inhibition. It will be important in future work to better identify and characterize the interrelationships among these.

