## Supplementary figures and images for "Gas1-Mediated Suppression of Hepatoblastoma Tumorigenesis"

### Supplemental Figure 1

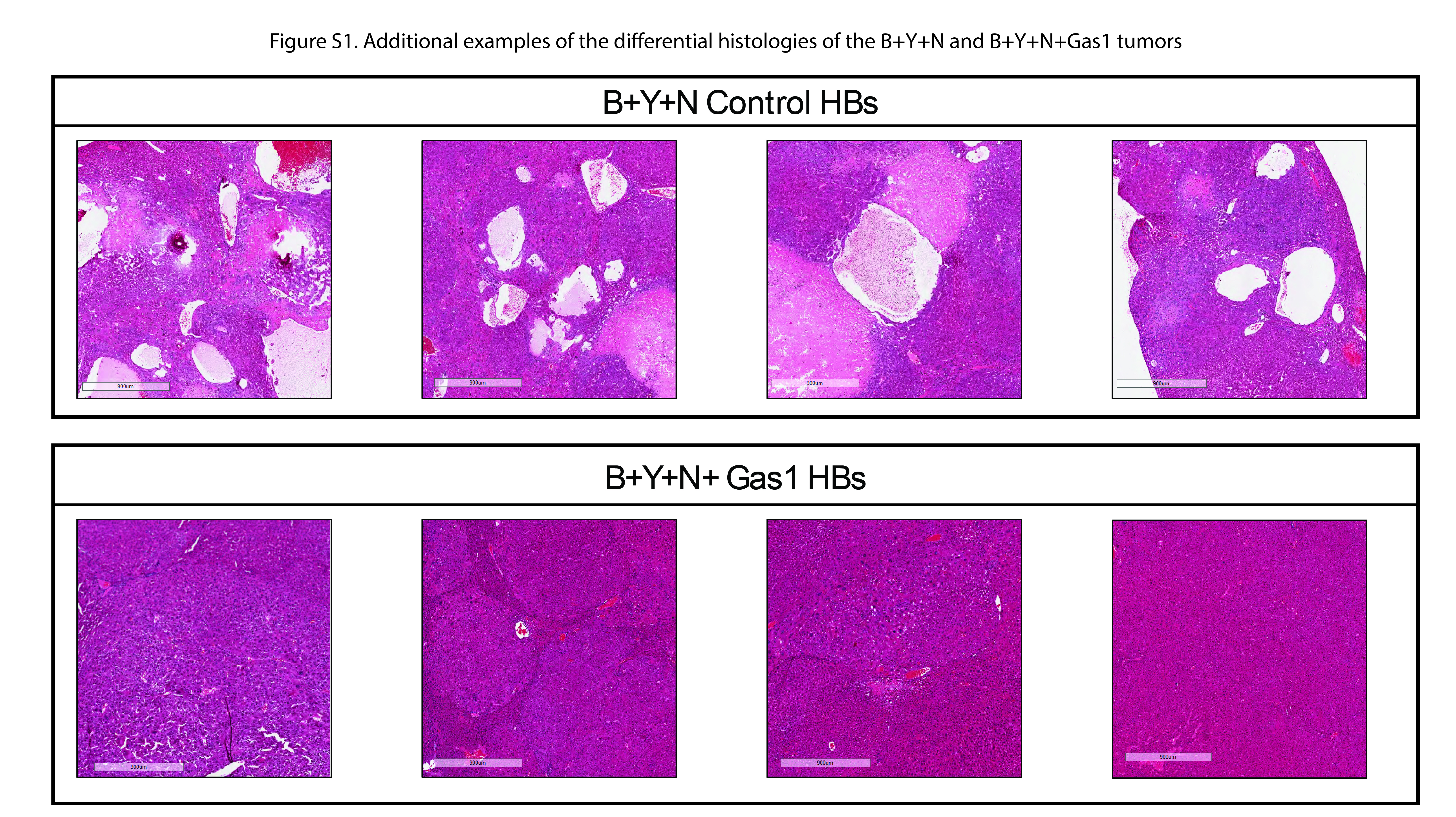
